# *Kupeantha yabassi* (Coffeeae- Rubiaceae), a new Critically Endangered shrub species of the Ebo Forest area, Littoral Region, Cameroon

**DOI:** 10.1101/2021.03.21.436301

**Authors:** Maria G. Alvarez-Aguirre, Martin Cheek, Bonaventure Sonké

## Abstract

A new species to science of evergreen forest shrub, *Kupeantha yabassi* (Coffeeae - Rubiaceae), is described, illustrated, mapped, and compared morphologically with the closely similar species *K. pentamera*. Restricted so far to a single site in evergreen lowland forest near the Ebo Forest, Yabassi, Littoral Region, Cameroon, this species is Critically Endangered using the IUCN 2012 standard due to habitat clearance driven mainly by agriculture, adding to the growing list of threatened species resulting from anthropogenic pressure on Cameroon forests. A revised key to the six species of *Kupeantha* is presented. Two distinct geographical and ecological species groupings within the genus are identified and discussed. Notes are given on other narrowly endemic and threatened species in the Ebo forest area, a threatened centre of diversity important for conservation in Littoral Region.

## Introduction

The new species reported in this paper was discovered as a result of a long-running survey of plants in Cameroon to support improved conservation management. The survey is led by botanists from the Royal Botanic Gardens, Kew and IRAD (Institute for Research in Agronomic Development)-National Herbarium of Cameroon, Yaoundé. This study has focussed on the Cross-Sanaga interval (Cheek *et al*. 2001, 2006) which contains the area with the highest plant species diversity per degree square in tropical Africa (Barthlott *et al*. 1996). The herbarium specimens collected in these surveys formed the foundations for a series of Conservation Checklists (see below). So far, over 100 new species and several new genera have been discovered and published as a result of these surveys, new protected areas have been recognised and the results of analysis are feeding into the Cameroon Important Plant Area programme (https://www.kew.org/science/our-science/projects/tropical-important-plant-areas-cameroon), based on the categories and criteria of Darbyshire *et al*. (2017).

During completion of a paper erecting the genus *Kupeantha* Cheek (Cheek *et al*. 2018a), it was noted that a specimen, *Leeuwenberg* 6400 (K), included in the protologue of *K. pentamera* (Robbr. & Sonké) Cheek (originally described as *Calycosiphonia pentamera* Robbr. & Sonké (Sonké *et al*. 2007), was geographically disjunct from all the other 38 specimens known of that species. Further investigation of this specimen, in connection with preparation of a Conservation Checklist of the Plants of the Ebo Forest, Littoral Region, has shown that the points of difference between this specimen and the other specimens of *Kupeantha pentamera* (see Table 1 below), are more than sufficient to warrant erection of a new species, the sixth in *Kupeantha*. We propose in this paper to provide evidence to test this hypothesis and to name this species *Kupeantha yabassi*.

**Table 1.** Main distinctions between *Kupeantha yabassi* and *K. pentamera*

| Character | *Kupeantha yabassi* | *Kupeantha pentamera* |
| --- | --- | --- |
| Leaf blade colour | Drying dark green | Drying black |
| Secondary nerves | Deeply impressed, giving a sub-bullate appearance to the leaf surface | Flush with surface, the leaf completely flat |
| Petiole length | 6 – 12 mm | (7 –) 9 – 13 mm |
| No. flowers per axil | (2 –) 3 (– 5) | 1 (– 2) |
| Lower calyculus dimensions | 1.2 – 1.5 x 1.2 – 1.5 mm | 0.5 – 1 × 0.5 – 1 mm |
| Upper calyculus dimensions | 2.2 – 3.5 x 1 – 1.5 mm | 1 – 1.5 × 1 – 2 mm |
| Fruit size (dried) | 10 – 13 x 11 – 14 mm | 17 – 25 x 13 – 16.5 mm |

*Kupeantha* Cheek (Coffeeae, Rubiaceae) is a recently described genus in the *Argocoffeopsis* clade which comprises also of *Argocoffeopsis* Lebrun and *Calycosiphonia* Pierre ex Robbr. (Cheek *et al*. 2018a). In that paper two new species were published (*Kupeantha ebo* M. Alvarez & Cheek and *Kupeantha kupensis* Cheek & Sonké, while the other three species had previously been described in either *Argocoffeopsis* or *Calycosiphonia* ((*Kupeantha fosimondi* (Tchiengué & Cheek) Cheek (Cheek & Tchiengue in Harvey *et al*. 2010), *Kupeantha pentamera* (Sonké & Robbr.) Cheek (Sonké *et al*. 2008), and *Kupeantha spathulata* (A.P.Davis & Sonké) Cheek (Davis & Sonké 2010)). *Kupeantha* is characterised by having supra-axillary buds, distal stem internodes drying dull black, proximal internodes with smooth, white spongy epidermis. The species all occur in Cameroon, with one of the species extending into Rio Muni of Equatorial Guinea. *Kupeantha* are found mainly in primary tropical forest from 100 to 2000 m altitude (Cheek *et al*. 2018a). It is possible that in future the range of the genus might be extended to adjacent Gabon, since over 1000 specimens of Rubiaceae were reported unidentified from that country by Sosef *et al*. (2005).

## Materials and Methods

Herbarium citations follow Index Herbariorum (Thiers *et al*. 2020). Specimens were viewed at BR, K, P, WAG, and YA. The National Herbarium of Cameroon, YA, was also searched for additional material of the new species, but without success. During the time that this paper was researched, it was not possible to obtain physical access to material at WAG (due to the transfer of WAG to Naturalis, Leiden, subsequent construction work, and covid-19 travel and access restrictions). However images for WAG specimens were studied at https://bioportal.naturalis.nl/?language=en and those from P at https://science.mnhn.fr/institution/mnhn/collection/p/item/search/form?lang=en_US. We also searched JStor Global Plants (2020) for additional material, and finally the Global Biodiversity Facility (GBIF, www.gbif.org accessed 23 Aug 2020).

Binomial authorities follow the International Plant Names Index (IPNI, 2020). The conservation assessment was made using the categories and criteria of IUCN (2012). Herbarium material was examined with a Leica Wild M8 dissecting binocular microscope fitted with an eyepiece graticule measuring in units of 0.025 mm at maximum magnification. The drawing was made with the same equipment using Leica 308700 camera lucida attachment. The terms and format of the description follow the conventions of Cheek *et al*. (2018a). The georeference for *Leeuwenberg* 6400 was obtained from the geographical metadata of the specimen using Google Earth (https://www.google.com/intl/en_uk/earth/versions/). The map was made using SimpleMappr (https://www.simplemappr.net).

## Results

### Taxonomic treatment

*Kupeantha yabassi* can be distinguished from *K. pentamera* which it is most closely similar to, and most easily confused with, using the diagnostic characters presented in Table 1, below. It can be distinguished from all other species of the genus using the key below.

Key to the species of *Kupeantha* (updated from cheek *et al*. 2018a)

1. Fruit obovoid; acumen spathulate ...................................................................... ***K. spathulata***

1. Fruit globose or ellipsoid; acumen with apex acute ............................................................... 2

2. 0.5 – 1.5m tall; tertiary nerves conspicuous, scalariform or forming fine reticulation on lower surface of leaf-blade; secondary nerves (9 –)10 – 13 on each side of midrib ............ 3

2. 2 – 5m tall; quaternary nerves absent or inconspicuous, scalariform or fine reticulation absent; secondary nerves <10 on each side of midrib ........................................................... 4

3. Leaves drying black, secondary nerves flush with adaxial surface; inflorescences 1(– 2) per axil; fruit 17 – 25 mm long .................................................................... ***K. pentamera***

3. Leaves drying green, secondary nerves deeply impressed on adaxial surface; inflorescences (2 –)3(– 5) per axil; fruit 10 – 13 mm long .......................................................... ***K. yabassi***

4. Fruits ellipsoid, ripening black, with a short stipe & rostrum. SW Region, Mt Kupe ............................................................................................................................. ***K. kupensis***

4. Fruit globose, ripening orange-red, stipe & rostrum absent or inconspicuous. SW Region,

Lebialem or Littoral Region, Ebo ......................................................................................... 5

5. Fruit 25 – 30 mm diameter. Lebialem Highlands of SW Region; 1300 – 1400 m alt ......................................................................................... ***K. fosimondi***

5. Fruit 10 – 15 mm diameter. Ebo Highlands of Littoral Region; 770 – 830 m alt ......... ***K. ebo***

### Kupeantha yabassi

*M*.*G*.*Alvarez & Cheek* **sp. nov**. Type: Cameroon, Littoral Region, 3 km east of km 21 of road Yabassi–Douala, 4° 19’ 12” N, 10° 04’ 09” E, fr, 17 Aug.1965, *A*.*J*.*M. Leeuwenberg* 6400 (holotype K000593314; isotypes WAG, BR, EA, LISC, MO, PRE, YA). (Fig. 1).

**Fig. 1.**
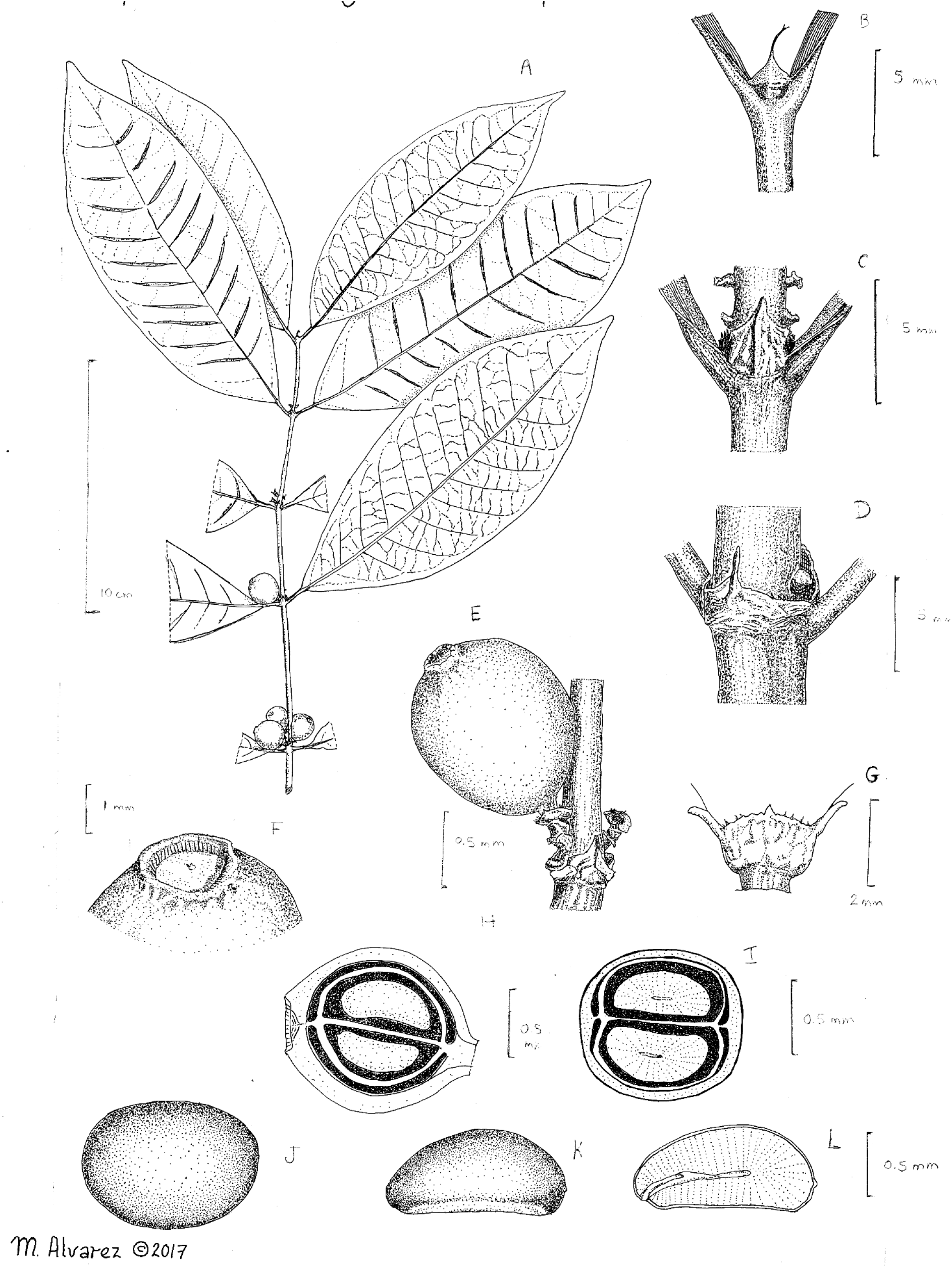
Kupeantha yabassi. **A** habit, fruiting branch; **B** detail of stipules at stem apex; **C & D** sheathing stipules and supra-axillary buds at subapical nodes; **E** fruit, showing calyculi; **F** fruit apex showing disc; **G** distal (upper) calyculus in young fruit; **H** fruit, longitudinal section showing the exocarp (dotted), mesocarp (white areas), cavities (dark areas) and at the centre, the seeds; **I** transverse section of a fruit, showing the exocarp (dotted), mesocarp (white areas), cavities (dark areas) and at the centre, endosperm with embedded embryo; **J** adaxial view of seed; **K** lateral view of seed; **L** longitudinal section of seed showing the endosperm (hatched) and part of embryo. **A-L** all drawn from A.J.M. *Leeuwenberg* 6400 by MARIA ALVAREZ.

*Evergreen shrub* 1.50 m tall, erect. Leafy branchlets with distal internodes drying black, black, terete, 2 – 3 mm in diameter, internodes 3.8 – 5.5 cm long, glabrous, smooth; epidermis of older branches becoming white, flaking and peeling. Stipules shortly sheathing, subcoriaceous, 2 – 3 x 1.75 – 4 (– 5) mm, keeled along midline, generally from the base, limb triangular, apex subulate, 0.5 – 2 mm long, detaching and falling with age, outer surface glabrous, inner surface with standard colleters (Fig. 1B-D).

*Leaves* opposite, equal; leaf–blades slightly bicoloured, drying dark green-brown above and grey-green below, papery, narrowly elliptic to oblong, 8 – 17 x 3 – 7 cm,; apex acuminate, 8

12 mm long; base attenuate to cuneate; midrib slightly raised above and prominent below; secondary nerves deeply impressed above (leaf surface approaching bullate), prominent below, 9 – 12 on each side of the midrib, ascending and uniting to make a looped intramarginal nerve (brochidodromous) 2 – 2.25 mm from the margin; midrib and secondary nerves drying black on abaxial surface; tertiary nerves scalariform to broadly reticulate on both sides of the leaf (Fig. 1A), glabrous above and below, domatia absent. Petioles drying black to dark brown, plano-convex, 0.6 – 1.1 cm long, 1 mm wide, the adaxial surface flat, glabrous. *Inflorescences* supra-axillary, inserted 2 – 4 mm above the axil, in consecutive nodes (Fig. 1A), (2 –) 3 (– 5) per axil, in opposite axils, 1-flowered, subsessile, peduncle 0.8–1.5 mm long. *Calyculi* 2, subsessile, cupuliform, glabrous; the first, proximal, calyculus, closer to the stem, often 4-lobed, 1.2 – 1.5 x 1.2 – 1.5 mm; the second, distal, calyculus, 2.2 – 3.5 x 1 – 1.5 mm, lobes 2, subulate, each 1 mm long (Fig. 1G). *Fruits* berry-like, orange at maturity, shortly ellipsoid to globose, 2 x 1.5 cm when fresh (field notes); 1.1 – 1.4 x 1.0 – 1.3 cm when dried (Fig. 1 E); subsessile, glabrous, exocarp drying hard, leathery, dark brown; disc ± circular, flat, 2 mm diam, sunk below calyx rim (Fig. 1F); calyx limbs absent. *Seeds* 2, apparently without seed coat (Fig. 1I), plano–convex, oblate in outline 9 – 12 x 6 – 10 x 2 – 6 mm. surface smooth with hilum elliptic, c. 1 mm long on ventral face (Fig. 1K), endosperm entirely cream, striated in longitudinal section, embryo about 0.5 mm long (Fig. 1L). Micropyle not detected.

## RECOGNITION

*Kupeantha yabassi* is an evergreen shrub, similar to *K. pentamera* (Sonké & Robbr.) Cheek, but differing in the fruits being smaller, 10 – 13 mm long (versus 17 – 25 mm), inflorescence more numerous, (2 –)3(– 5) per axil (versus 1(– 2), upper calyculus 2.2–3.5 mm long (versus 1 – 1.5 mm long), leaf-blade drying green, sub-bullate due to the impressed secondary nerves (not black, surface flat). Additional characters distinguishing *Kupeantha yabassi* from *K. pentamera* are given in Table 1.

## DISTRIBUTION

Endemic to Cameroon (Map 1). Only known from a single collection made in forest near the Ebo Forest in the Littoral Region.

**Map 1.**
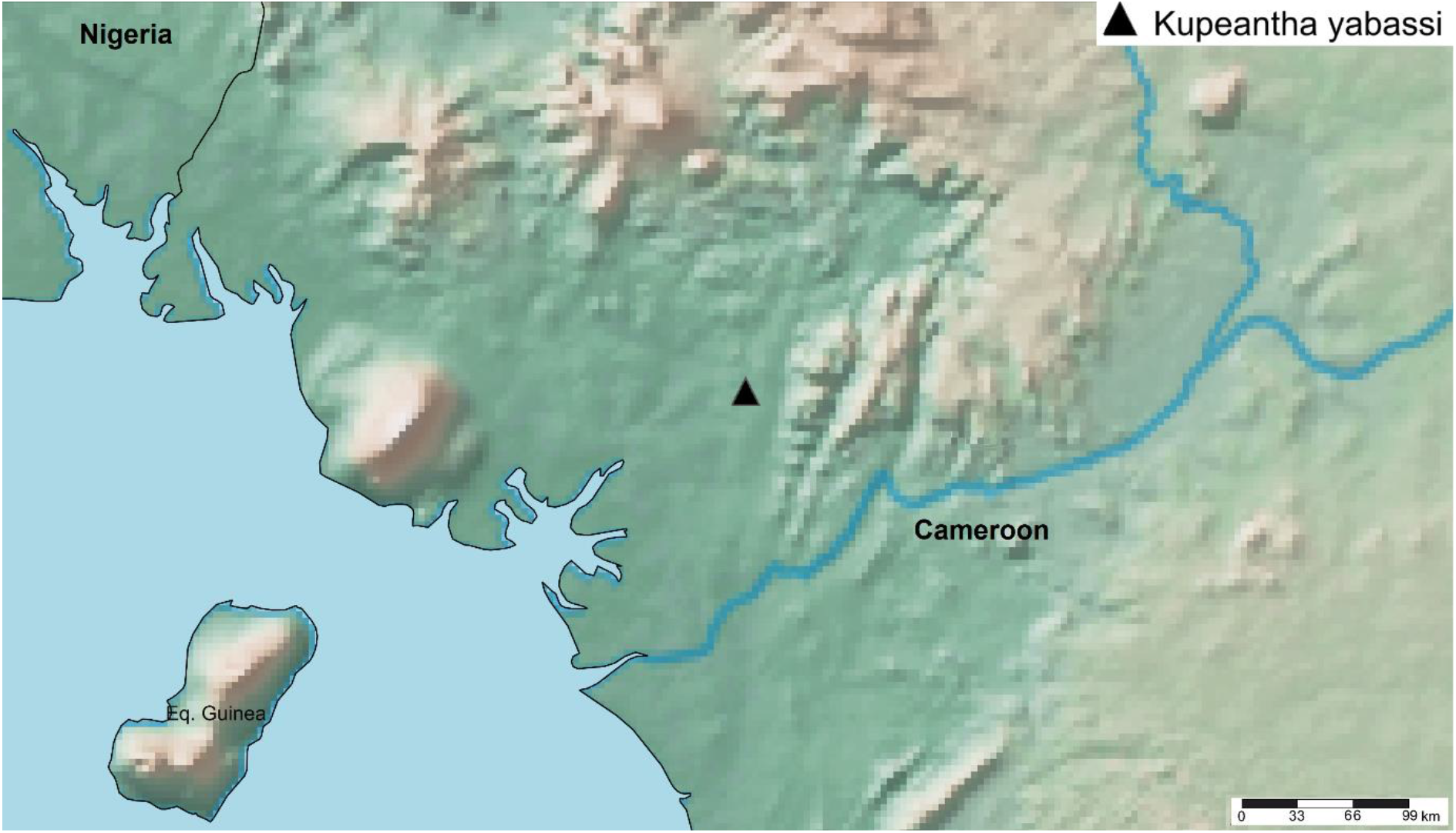
Known global distribution of *Kupeantha yabassi*.

## HABITAT

Secondary forest, along the Yabassi-Douala road, with *Lophira alata*,Banks ex C.F.Gaertn., *Coula edulis* Baill. and *Sacoglottis gabonensis* (Baill.) Urb., 50 – 100 m altitude. The presence of *Lophira alata* and *Sacoglottis gabonensis* in rainforest within c. 50 – 100 km from the coast in Cameroon, indicates that the forest has been cultivated in the past (Letouzey 1957, 1960, 1968, 1985). At Ebo the geology is ancient, highly weathered basement complex, with some ferralitic areas in foothill areas which are inland, c. 100 km from the coast. The wet season (successive months with cumulative rainfall >100 mm) falls between March and November and is colder than the dry season (Abwe & Morgan 2008).

## CONSERVATION STATUS

*Kupeantha yabassi* is known from only a single site along the road parallel to the western boundary of the Ebo forest. Since 2006, botanical surveys have been mounted almost annually, at different seasons, over many parts of the formerly proposed National Park of Ebo. About 3000 botanical herbarium specimens have been collected, but this species has been seen not yet been seen in the c. 2000 km^2^ of the Ebo Forest. However, much of this area, especially the western edge, has not yet been surveyed for plants. While it is likely that the species will be found at additional site likely including the Ebo Forest, there is no doubt that it is genuinely range-restricted. Botanical surveys and other plant studies for conservation management in forest areas north, west and east of Ebo resulting in thousands of specimens being collected and identified have failed to find any additional specimens of this species (Cheek *et al*. 1996; Cable & Cheek 1998; Cheek *et al*. 2000; Maisels *et al*. 2000, Harvey *et al*. 2004; Cheek *et al*. 2004; Cheek *et al*. 2010; Harvey *et al*. 2010; Cheek *et al*. 2011). It is possible that the species is truly localised as are all other species of the genus west of the Sanaga (see discussion below). *Kupeantha yabassi* may well be unique to the Ebo Forest area, as are also, on current evidence, at least five other species (see discussion, below). The area of occupation of *Kupeantha yabassi* is estimated as 4 km^2^ using the IUCN preferred cell-size. The extent of occurrence is the same. In February 2020 it was discovered that moves were in place to convert the forest into two logging concessions (e.g. https://www.globalwildlife.org/blog/ebo-forest-a-stronghold-for-cameroons-wildlife/ and https://blog.resourceshark.com/cameroon-approves-logging-concession-that-will-destroy-ebo-forest-gorilla-habitat/ both accessed 19 Sept. 2020). Such logging would result in timber extraction that would open up the canopy and remove the intact habitat in which *Kupeantha yabassi* is thought to grow. Additionally, slash and burn agriculture often follows logging trails and would negatively impact the population of this species. Fortunately the logging concession was suspended in August 2020 due to representations to the President of Cameroon on the global importance of the biodiversity of Ebo (https://www.businesswire.com/news/home/20200817005135/en/Relief-in-the-Forest-Cameroonian-Government-Backtracks-on-the-Ebo-Forest accessed 19 Sept. 2020). However, the forest habitat of this species remains unprotected and threats of logging and conversion of the habitat to plantations remain. *Kupeantha yabassi* is therefore here assessed, on the basis of the range size given and threats stated as CR B1+2ab(iii), that is Critically Endangered.

## PHENOLOGY

Flowering unknown; fruiting in August.

## ETYMOLOGY

*Kupeantha yabassi* named for the administrative centre nearest to the point where the specimen was collected.

## VERNACULAR NAMES

No vernacular names or uses are recorded.

## NOTES

Only known from the single collection cited. At Kew the specimen was annotated “indet. not matched in *Tricalysia* etc.”, subsequently, loaned to BR in 1983 it was annotated by Robbrecht in 1986 as “unknown to me, I don’t believe it belongs to *Tricalysia*”

## Discussion

### Groupings within *Kupeantha*

The former placement of the only known specimen of *Kupeantha yabassi* in *K. pentamera* was understandable due to their morphological similarity. Both species have leaves of similar shape and size, and uniquely in the genus have tertiary nerves which are visible (they are inconspicuous in other species of the genus). They also share fruits of similar shape.

The six species of *Kupeantha* can be divided into two groups based on ecology and geography. The groups differ from each other also in sympatry and extent of occurrence.

1) East of the Sanaga river, the only two species documented are *Kupeantha spathulata* and *K. pentamera*. Both species are low altitude, predominantly occurring below 800 m (although the last has been recorded as high as 900 m). Both are known from numerous specimens 21 and 37 respectively, with numerous locations and large extents of occurrence (8105 km^2^, and 52 km^2^ respectively). The two species are often sympatric, indeed have been collected in sequential number series at several of their common locations, in South Region, Cameroon: e.g. Bibondi, 24 Jan. 2005, *Sonké & Nguembou* 3783 (*K. pentamera*) and 3784 (*K. spathulata*). Again, at 3 km NNW Ngoyang, 20 Sept. 2005, *Sonké & Djuikouo 4060 (K. spathulata*), and 4059 (*K. pentamera*). Similarly, at 2 km NW Mbikililiki, 19 Jan. 2006, *Sonké & Djuikouo* 4286 (*K. pentamera*) and 4285 (*K. spathulata*). These specimens are cited in the protologues of the two species (Sonké *et al*. 2008 and Davis & Sonké 2010), they derive from a series of Rubiaceae-focussed surveys that resulted in numerous other discoveries of new Rubiaceae species to science, e.g., Sonké (2005, 2006 & 2008).

2) West of the Sanaga River, the four species known, *Kupeantha kupensis, K. fosimondi, K*.*ebo and K. yabassii* are all upland, cloud forest species, occurring in the 800 – 2000 m altitudinal range (However, *Kupeantha ebo* occurs in the range 770 – 832 m). None are sympatric but allopatric, all are separated from each other by tens of kilometres. Even in the Ebo forest area, *Kupeantha ebo* and *K. yabassi* are physically separated by 20 km. These four species all have much smaller extents of occurrence (c. 8 km^2^ in *K. kupensis*) and are much rarer and more localised (and so more threatened) than those species east of the Sanaga.

Despite the separation of these two groups, there is no evidence that they are separated phylogenetically. The two species East of the Sanaga are in fact so dissimilar that they were formerly placed apart from each other in separate genera (*Calycosiphonia* and *Argocoffeopsis*, Sonké *et al*. 2008 and Davis & Sonké 2010 respectively).

### *Kupeantha yabassi*: its range and other endemic species in the Ebo Forest area

Abwe & Morgan, (2008) and Cheek *et al*. (2018b) characterise the Ebo forest which is adjacent to the location of *Kupeantha yabassi*, and give overviews of habitats, species and importance for conservation. Fifty-two globally threatened plant species are currently listed from Ebo on the IUCN Red List website and the number is set to rise rapidly as more of Cameroon’s rare species are assessed for their conservation status as part of the Cameroon TIPAs programme. The discovery of a new species to science near the Ebo forest is not unusual. Numerous new species have been published from Ebo in recent years. Examples of other species that, like *Kupeantha yabassi*, appear to be strictly endemic to the Ebo area on current evidence are: *Ardisia ebo* Cheek (Cheek & Xanthos, 2012), *Crateranthus cameroonensis* Cheek & Prance (Prance & Jongkind, 2015), Gilbertiodendron *ebo* Burgt & Mackinder (van der Burgt *et al*. 2015), *Inversodicraea ebo* Cheek (Cheek *et al*. 2017), *Kupeantha ebo* M.Alvarez & Cheek (Cheek *et al*. 2018a), *Palisota ebo* Cheek (Cheek *et al*. 2018b) and *Pseudohydrosme ebo* Cheek (Cheek *et al*. 2021).

Further species described from Ebo have also been found further west, in the Cameroon Highlands, particularly at Mt Kupe and the Bakossi Mts (Cheek *et al*. 2004). Examples are *Myrianthus fosi* Cheek (Cheek & Osborne, in Harvey *et al*. 2010), *Salacia nigra* Cheek (Gosline & Cheek, 2014), *Talbotiella ebo* Mackinder & Wieringa (Mackinder *et al*. 2010)

Additionally, several species formerly thought endemic to Mt Kupe and the Bakossi Mts have subsequently been found at Ebo, e.g. *Coffea montekupensis* Stoff. (Stoffelen *et al*. 1997), *Costus kupensis* Maas & H. Maas *(Maas-van der Kamer et al. 2016), Microcos magnifica* Cheek (Cheek, 2017), and *Uvariopsis submontana* Kenfack, Gosline & Gereau (Kenfack *et al*. 2003). It is considered likely that additional Kupe species may yet be found at Ebo such as *Brachystephanus kupeensis* I.Darbysh. (Champluvier & Darbyshire, 2009), *Impatiens frithii* Cheek (Cheek & Csiba 2002) since new discoveries are still frequently being made in the Ebo Forest area. Therefore, it is possible that *Kupeantha yabassi* might yet also be found in the Cameroon highlands, e.g. at Mt Kupe. However, this is thought to be only a relatively small possibility given the high level of survey effort at Mt Kupe: if it occurred there it is highly likely that it would have been recorded already.

Cameroon has been highlighted as the top country in tropical Africa for plant species diversity per degree square, and has high levels of endemism (Lachenaud *et al*. 2013; Onana 2011, Onana & Cheek 2011). The inventory of its Flora is far from completed, and the few remaining sizable areas of intact forest such as Ebo Forest, in the Littoral Region, are seriously threatened by clearance. Unfortunately, there is an increasing pressure to convert critical areas for biodiversity into oil palm plantations, and logging concessions (Mahmoud *et al*. 2019). As stated in the introduction, efforts are being made to delimit the highest priority areas in Cameroon for plant conservation as Tropical Important Plant Areas (TIPAs) (Darbyshire *et al*. 2017) to avoid the extinction of narrowly endemic species (Cheek et al 2018a; Mahmoud *et al*. 2019). However, the pressure for forest and wildlife resources is a stronger driver in the rapid decrease of primary forests and biodiversity. This paper contributes to documenting the rich flora of Cameroon and helps to highlight areas for conservation.

Such discoveries as this new species also underline the urgency for making such further discoveries while it is still possible since in all but one of the cases given, the range extension resulted from discovery of a new species for science with a narrow geographic range and/or very few individuals, and which face threats to their natural habitat, putting these species at high risk of extinction.

About 2000 new species of vascular plant have been discovered each year for the last decade or more. Until species are known to science, they cannot be assessed for their conservation status and the possibility of protecting them is reduced (Cheek *et al*. 2020). Documented extinctions of plant species are increasing, e.g. *Oxygyne triandra* Schltr. and *Afrothismia pachyantha* Schltr. of South West Region, Cameroon are now known to be globally extinct (Cheek & Williams, 1999, Cheek *et al*. 2018c, Cheek *et al*. 2019). In some cases species appear to be extinct even before they are known to science, such as *Vepris bali* Cheek, also from the Cross-Sanaga interval in Cameroon (Cheek *et al*. 2018d) and elsewhere, *Nepenthes maximoides* Cheek (*King & Cheek, 2020*). Most of the >800 Cameroonian species in the Red Data Book for the plants of Cameroon are threatened with extinction due to habitat clearance or degradation, especially of forest for small-holder and plantation agriculture following logging (Onana & Cheek, 2011). Efforts are now being made to delimit the highest priority areas in Cameroon for plant conservation as Tropical Important Plant Areas (TIPAs) using the revised IPA criteria set out in Darbyshire *et al*. (2017). This is intended to help avoid the global extinction of additional endemic species such as *Kupeantha yabassi* which we hope will be included in the proposed Ebo Forest IPA.

With only one locality known, *Kupeantha yabassi* represents another narrowly endemic Cameroonian species threatened with extinction due to deforestation for oil palm plantations, small-scale agriculture, mining and logging (Onana & Cheek 2011, Cheek *et al*. 2018a).

## Acknowledgements

This paper was completed as part of the Cameroon TIPAs (Tropical Important Plant Areas) project at RBG, Kew, which is supported by Players of People’s Postcode Lottery. Ekwoge Abwe and Bethan Morgan and their team at the Ebo Forest programme are thanked hugely for expediting our botanical surveys in the Ebo forest of Cameroon over several years which allowed us to give context about the Ebo forest in this paper.

The heads of IRAD (Institute of Research in Agronomic Development)-National Herbarium of Cameroon, Yaoundé, successively Jean-Michel Onana, Florence Ngo Ngwe, Eric Nana and Jean Betti Lagarde, are thanked for co-ordinating the co-operation with the Royal Botanic Gardens, Kew.

The authors would like to thank two anonymous reviewers for comments on an earlier version of this manuscript.

